# Plant carbohydrate depletion impairs water relations and spreads via ectomycorrhizal networks

**DOI:** 10.1101/2020.08.03.234823

**Authors:** Gerard Sapes, Patrick Demaree, Ylva Lekberg, Anna Sala

## Abstract

Carbon and water relations are fundamental to plant life and strongly interact. Under drought, the ability of plants to assimilate carbon is reduced, which increases their consumption of stored labile carbon in the form of non-structural carbohydrates (NSC). Stored NSC depletion may impair plant water relations, but mechanisms are not clear, and we do not know if its effects are independent of water deficit. If so, carbon costs of fungal symbionts could also indirectly influence plant drought tolerance through stored NSC depletion. We connected well-watered *Pinus ponderosa* seedling pairs via ectomycorrhizal (EM) networks where one seedling was shaded and the other experienced full light and compared responses to seedling pairs in the light. We measured plant water relations and traced carbon movements using ^13^CO_2_ to explore the mechanisms linking stored NSC to water relations, and to identify potential tradeoffs between the ability to endure low water potentials and maintaining EM fungi under carbon-limiting conditions. Even in the absence of drought, mild NSC depletion impaired osmoregulation capacity and turgor maintenance, a critical strategy to tolerate drought. This demonstrates that NSC storage influences plant water relations independently of plant water status. We also found that EM networks propagated NSC depletion and its negative effects on water relations from carbon stressed hosts to non-stressed hosts. These results highlight carbon allocation tradeoffs between supporting fungal symbionts and retaining water via stored NSC and have implications for biotic interactions and forest drought responses.

**Significance Statement:** The potential effects of future drought on global carbon cycles, vegetation-climate feedbacks, species distributions and their ecological impacts, urgently call for a clear understanding of factors influencing vegetation tolerance to drought. Key to this is the understanding of mechanisms and processes by which plants tolerate drought and how prevalent plant-fungal interactions may influence these processes.

We demonstrate that even mild depletion of plant non-structural carbohydrate (NSC) storage readily decreases plant water retention capacity, therefore decreasing tolerance to drought. Because plant-fungal interactions depend on NSC exchange, plants face carbon-allocation tradeoffs between maintaining drought tolerance and feeding fungal symbionts. The impacts of these tradeoffs extend across plants connected via ectomycorrhizal networks as fungi propagate NSC depletion from NSC-limited plants to non-stressed individuals.

## Introduction

Under future climate, more intense and frequent droughts are expected to increase forest mortality and to impact forest ecosystems and global carbon cycles ^1^. Predicting the extent of these impacts requires a full understanding of the mechanisms and processes associated with drought-induced mortality (DIM) ^2,3^. Under drought, plants ultimately die from desiccation because water supply does not keep pace with water loss. Desiccation and death are generally associated with failure of the vascular system to transport water due to embolism formation (hydraulic failure, Adams *et al.* 2017). Reduced photosynthesis under prolonged drought eventually leads to the depletion of stored non-structural carbohydrates (NSC) and carbon limitation, a process thought to contribute to desiccation by decreasing the plant’s ability to retain and move water ^5–8^. Hydraulic failure is thought to be the main driver of DIM with NSC depletion playing a secondary, interacting role. However, it is not known whether NSC depletion *per se* impairs plant water relations because its effects have not been experimentally isolated from those caused by drought (e.g., reduced water transport) (Hartmann and Trumbore 2016, Adams et al. 2017). Yet, NSC depletion may be more relevant than currently thought as the wide array of physiological processes that demand NSC may limit carbon allocation to storage and its positive effects on water relations. For instance, in addition to metabolic costs, many trees incur significant carbon costs to maintain symbiotic partnerships with ectomycorrhizal (EM) fungi ^11^. Consequently, plants may face carbon allocation tradeoffs between sustaining symbionts and maintaining water relations via NSC storage under carbon-limiting conditions. Whether these tradeoffs exist has not been tested despite their potential implications for forest mortality under global change. In this study, we examine the independent effects of stored NSC for plant water relations and assess potential tradeoffs between maintaining water relations and sustaining EM symbionts when carbon becomes limiting.

Stored NSC have been classically understood as a carbon safeguard for growth and energy. However, NSC may play an important role in the movement and retention of water ^5^ including lowering tissue osmotic potential to increase tolerance to low water potentials under drought ^12^. While the osmotically active soluble fraction of NSC plays a direct osmotic role, starch could also play an indirect role as a source of osmotically active organic compounds and energy ^13^. Under carbon-limiting conditions of any kind, NSC-consuming processes such as respiration, defense production, and maintenance of ectomycorrhizae rely on the same pool of stored NSC potentially involved in osmoregulation ^14^. Consistently, complete consumption of stored NSC is rare under natural and experimental conditions, and plants rarely deplete their stored NSC below ca. 40% of their seasonal maximums ^4,15^. While NSC have been associated with maintenance of hydraulic conductivity, capacitance, and turgor ^9,16,17^, it is unclear how much of the total stored NSC pool plants must conserve to maintain these functions, even under non-drought conditions ^18^. If a significant fraction of stored NSC must remain unconsumed to maintain the capacity to move and retain water, we may have underestimated the role of NSC depletion in DIM, which could have important implications for modelling and prediction of tree mortality.

In temperate coniferous and boreal ecosystems, virtually all trees and seedlings form associations with EM fungi and establish underground EM networks through which resources can be exchanged ^19–21^. These networks have been shown to enhance seedling establishment and increase ecosystem-level nutrient cycling ^22,23^. Plant hosts allocate up to 30% of their assimilated carbon to EM fungi ^11,24^ in exchange for nutrients. As such, EM fungi could indirectly influence tolerance to drought via network-mediated carbon redistribution and host NSC depletion under carbon-limiting conditions. For instance, if stochastic or microsite effects (e.g., herbivory, shading, microsite low moisture) cause carbon deficit and NSC depletion on some plant hosts but not others within the network, NSC-depleted hosts could decrease carbon supply to their associated EM fungi and trigger fungal carbon deficit. Non-NSC-depleted hosts may then offset the deficit of the EM network with their own stored NSC, a response that could indirectly compromise their own water relations especially if they cannot regulate carbon flow to EM fungi. There is debate on whether carbon can be transferred among connected hosts via EM networks ^20,25,26^ or whether NSC simply stays within the EM fungi ^27^. In either case, network-mediated carbon redistribution may influence how individual plants and entire forest stands respond to drought. Whether and under what circumstances fungal carbon demand and redistribution compromise allocation to plant NSC storage and water relations is unknown but deserves attention.

Here, we used an experimental design that depletes plant NSC pools by combining shading treatments and EM networks among well-watered ponderosa pine seedlings (*Pinus ponderosa* Douglas ex C. Lawson). This design allowed us to unveil the mechanisms (and their importance) linking stored NSC and plant water relations without the confounding effects of water deficit, as well as the carbon dynamics between ectomycorrhizal fungi and their plant hosts under carbon-limiting conditions. Our experiment involved seedlings as they often determine forest distributions through establishment ^28^. Also, because of their small size and little storage capacity, seedlings may be especially susceptible to carbon allocation tradeoffs between maintaining water relations and EM fungi. Specifically, we asked 1) does stored NSC depletion affect host water relations even without water deficit?; and 2) do ectomycorrhizal networks extend the effects of NSC depletion on plant water relations to non-stressed neighbors? We found that, even in the absence of drought, mild NSC depletion indirectly decreased plant tolerance to drought by impairing osmoregulation capacity and turgor maintenance. These results unambiguously link NSC storage to plant water relations. We also found that EM networks propagated NSC depletion and impaired critical drought tolerance strategies in non-stressed hosts. This demonstrates the existence of resource-allocation tradeoffs between supporting fungal symbionts and retaining water via stored NSC and suggests that EM networks can affect tree and possibly entire forest tolerance to drought.

## Results

### NSC decreased in LD and D seedlings

Shading caused a 72% reduction in whole-plant NSC concentrations in dark (D) relative to light (LL) seedlings (p < 0.001). Light seedlings paired with dark (LD) plants also experienced NSC depletion (45% reduction) and showed intermediate whole-plant NSC pools relative to LL and D seedlings (p < 0.001, and p = 0.009, respectively) (Fig. 2). When broken down to each NSC compound, we found that starch was higher in LL seedlings (p < 0.001) relative to LD and D seedlings. LD seedlings had intermediate starch levels although only marginally higher than D seedlings (p = 0.079). Both sucrose and glucose + fructose were lower in D seedlings than in LL and LD seedlings (sucrose: LL: p < 0.001, LD: p = 0.040; glucose and fructose: LL: p < 0.001, LD: p = 0.018). However, no differences were detected between LL and LD seedlings. Patterns within each organ were consistent with those observed at the whole-plant level (Figure S6).

### Labeled carbon stayed within ectomycorrhizal fungi

Labeled LL and LD seedlings showed significant increases in ^13^C/^12^C ratios (*δ*^13^C) across all organs relative to pre-labeling base-line values (Fig. 3, Figure S7) while non-labeled seedlings stayed at base-line levels. This indicates that only labeled seedlings incorporated the C-isotope and that there was no contamination during the labeling process. In labeled seedlings, ^13^C progressively declined from needles to roots (p < 0.001 in all pairwise contrasts across seedling types and organs, Fig. 3, Figure S7: Panels a-c). High levels of ^13^C were also present in fungal tissue associated with labeled LL and LD seedlings (Fig. 3, Figure S8: Panel a). To a lesser extent, ^13^C was also present in fungal tissue associated with non-labeled LL plants paired with labeled LL plants (Fig. 3, Figure S8: Panel b). ^13^C in fungi associated with non-labeled LL plants confirmed the existence of a mycorrhizal network between seedlings at harvesting (Figure S1 and S2). However, ^13^C was not detected in colonized root tips of non-labeled D plants paired with labeled LD seedlings (Fig. 3, Figure S8: Panel b). In all cases, ^13^C levels within non-labeled seedlings were similar to base-line values regardless of organ or seedling type (Fig. 3, Figure S7: Panels d-f).

### LD and D seedlings exhibited lower turgor

Hydraulic conductivity, which describes the capacity to transport water, did not significantly differ among seedling types in any organ (Figure S9). However, osmotic potentials, which describe the ability to retain cell water at low water potentials, as is typical under drought, were up to ca. 1.5 MPa lower (i.e., higher water retention capacity) in LL seedlings than in LD and D seedlings (Fig. 4a-c). These differences were observed in needles (p < 0.001 in all pairwise contrasts), stems (D: p = 0.002, LD: p < 0.001), and roots (D: p < 0.001, LD: p = 0.001). High osmotic potentials were associated with low pressure potentials (i.e., turgor) in D and LD needles (D: p < 0.001, LD: p < 0.001, Fig. 4d) and stems (D: p = 0.012, LD: p < 0.001, Fig. 4e). Pressure potentials in LD and D stems were low enough to bring stems to turgor loss (i.e., pressure potential lower than 0). As a result, we observed lower stem relative water content in NSC-depleted seedlings than in non-depleted plants (D: p < 0.001, LD: p = 0.047, Figure S10).

### NSC depletion impaired water relations

NSC depletion in needles was associated with less negative (higher) needle osmotic potentials (p < 0.001, R = −0.83, Table S2, Fig. 5a). This relationship was linear when NSC depletion was large and plateaued when NSC were ca. 20% depleted relative to non-depleted seedlings. The pressure potential (necessary for positive turgor) mirrored this pattern (p < 0.001, R = 0.64, Table S2, Fig. 5b) and plateaued at 1.5 MPa when NSC was ca. 20% NSC depleted.

Stored NSC depletion also influenced the relationship between needle pressure potential and needle water potential (WP) (p < 0.001, R = 0.78, Fig. 6; Table S2). Plants with greater NSC depletion showed lower pressure at any given WP and higher WP at turgor loss (NSC deviation from controls: p < 0.001, Fig. 6). Additionally, plants with greater NSC depletion lost more turgor per unit decline in WP than less NSC-depleted plants with (Leaf WP x NSC deviation from controls: p = 0.006, Fig. 6).

## Discussion

It has long been known that plants exchange water (lost through stomata) for carbon (assimilated from atmospheric CO_2_). Our results demonstrate that plants also need stored labile carbon in the form of non-structural carbohydrates (NSC) to retain water. We asked 1) whether stored NSC depletion affect host water relations even without water deficit and 2) whether ectomycorrhizal networks extend the effects of NSC depletion on plant water relations to non-stressed neighbors. We found that NSC depletion impaired turgor maintenance - a critical drought tolerance strategy-even in well-watered plants. We also found that carbon-limiting conditions can result in a tradeoff between supporting symbionts and retaining water via stored NSC, and that this effect can be propagated by the EM network to otherwise non-stressed plants. During periods such as drought, when carbon supply via photosynthesis is limited, carbon allocation tradeoffs may be exacerbated by EM fungi and influence plant survival.

### Depletion of stored NSC impairs a critical drought tolerance strategy

Despite being well-watered, seedlings that experienced stored NSC depletion below 20% of non-NSC-depleted plants experienced declines in turgor and osmoregulation capacity (Figs. 4–6). Leaf osmotic potentials shifted from values typical of drought-tolerant temperate conifers (−2.3 MPa) to values greater than those found in drought-vulnerable croplands (−1.3 MPa) (Bartlett *et al.* 2012) (Fig. 4), which led to turgor loss (Fig. 6). These shifts were consistent with declines in total NSC concentrations but were not always reflected by declines in the osmotically active fraction of NSC (glucose, fructose, and sucrose). In fact, shifts in osmotic potential were mainly associated with declines in osmotically inactive starch. These results suggest that the effects of NSC depletion on osmoregulation are largely indirect, perhaps via the synthesis of osmotically active molecules other than soluble sugars (e.g., sugar alcohols, proteins, amino acids) ^29–31^. Regardless of the specific mechanisms, however, our results show that stored NSC depletion impairs a critical drought tolerance strategy in plants. The link between stored NSC and maintenance of osmotic and pressure potentials could explain previous findings that NSC-depleted plants experience lower phloem turgor under well-watered conditions ^9^, lower water content at any degree of water deficit ^6^, and reach lower stem WP and die at faster rates than NSC-enriched plants when exposed to similar water deficit ^16^. Based on our results, and given that osmotic potentials decrease from moist to dry biomes to compensate for low water potentials typical of dry regions ^12^, drought-tolerant plants from dry biomes might require greater NSC pools to maintain lower osmotic potentials than those from moist biomes and be more prone to experience carbon-allocation tradeoffs.

Drought impairs plant water transport and depletes NSC, and both have negative effects on plant water relations. Therefore, it is not surprising that both hydraulic failure and NSC depletion are often associated with drought-induced mortality (DIM) ^4,16^. However, despite an increasing interest in the potential role of stored NSC on plant water relations ^5,9,10^, evidence that NSC affects plant water relations independent of drought has been lacking. The negative impact of NSC depletion on osmoregulation capacity and turgor maintenance we show here provides such evidence. Our findings are consistent with the observation that plants prevent NSC depletion below certain thresholds at all costs ^15,32^, as well as observed increases in stomatal conductance in response to increases in osmotic potential ^33^. Together, these findings suggest that NSC depletion may further decrease drought tolerance by preventing total stomatal closure under drought. While still hypothetical, this could explain sudden increases in stomatal conductance at low water potentials ^32^, accelerated water loss ^6^ and accelerated declines in WP observed in drought-stressed, NSC-depleted plants ^16^ leading to early hydraulic failure and death. Our results provide support for a greater focus on plant water balance (turgor maintenance) in our efforts to understand interactions between water and carbon to better monitor and predict DIM ^6,7,34^. Mechanistic models that consider turgor loss may achieve higher species-specific predictive capacity because turgor loss marks the onset of many processes that lead to plant death. Additionally, turgor loss integrates many processes, including regulation of water transport, water loss and water retention. Additional research is needed to understand the specific mechanisms by which NSC depletion impairs osmoregulation across plant types, drought strategies, and environments and how species-specific carbon allocation strategies, including to storage and osmoregulation, influence tolerance to drought.

### Ectomycorrhizal networks can exacerbate NSC depletion and carbon allocation tradeoffs

We hypothesized that shaded seedlings (D) would indirectly increase carbon demand on connected, non-shaded seedlings (LD) due to a disruption of carbon allocation to EM fungi from D seedlings. Our results supported this hypothesis: LD seedlings became NSC-depleted relative to LL seedlings (Fig. 2), indicating that carbon demand in LD seedlings exceeded supply. Additional measurements confirmed that assimilation did not decrease in LD relative to LL seedlings (i.e. supply did not decrease; Figure S5). Likewise, there were no differences in growth and respiration between LD and LL seedlings (Figure S5). Clearly, stored carbon was lost in LD seedlings during the three weeks of shading, but it did not go to any other sink we could measure, leaving the EM network as the most likely sink. However, we did not find more ^13^C in the EM network in the LD pair than the LL pair, and the fungal tissue closest to the D seedling was not enriched, which is opposite to what we expected if the LD seedling bore a greater cost for supporting the network. This is not because the EM network connecting the seedlings was lacking. We confirmed the presence of the network both visually (Figures S1 and S2) and isotopically as ^13^C travelled from labeled seedlings to fungal tissue connected to neighboring, unlabeled seedlings (Fig 3). We believe that the most likely explanation is that, by the time labeling occurred, the LD seedling had either reduced carbon allocation to the EM network as costs exceeded benefits ^24,35,36^, or the EM fungi closest to the starved D seedling had died or become dormant, reducing the overall carbon demand on the LD seedling. While we cannot unambiguously attribute the missing carbon in LD seedlings to increased allocation to EM fungi (and possibly D seedlings ^20^), it does not change the fact that the EM network propagated NSC depletion to a non-stressed plant and that this had consequences for drought tolerance strategies. Thus, although EM fungi may increase survival by hydraulic lift ^19^, they may also reduce plant tolerance to drought ^37^ by contributing to the carbon allocation tradeoff between supporting symbionts and retaining water via stored NSC.

Carbon costs of sustaining EM may increase when plant or fungal carbon demand increases ^38^ and during periods of carbon limitation such as prolonged drought ^39^. Costs of sustaining EM networks might also increase substantially when some individuals connected to the network die (e.g. fire, drought, disease, and insect induced mortality) and the surviving trees must subsidize carbon supply to EM until fungal biomass re-adjusts to new levels of carbon supply. In these situations, EM networks might increase NSC depletion rates in hosts, hinder their osmoregulation capacity, and hasten desiccation rates. It is possible that our pairwise approach overestimates costs to individual seedlings and that EM network-driven NSC depletion is less substantial in complex networks of highly interconnected plants where increased costs are subsidized by several hosts ^40,41^. As such, EM networks may even enhance community resilience to drought, although this will likely depend on the proportion of stressed hosts within the network. The carbon costs of EM-networks under carbon-limiting conditions could also be offset over time by positive effects on overall performance or fitness that occur at times when carbon is not limiting (e.g., enhanced growth, photosynthetic rate, nutrient acquisition) ^42,43^. Studying all the factors that influence costs and benefits of EM networks at different spatiotemporal scales is critical to predict when and where they might have positive or negative effects under future climates ^44^. While studies often assess costs and benefits of mycorrhizal associations based on whether they increase or decrease plant growth ^45^, growth is only one of several functions requiring NSC and may not reflect costs and benefits for other vital plant functions, especially under stressful conditions that inhibit growth. Using a carbon balance approach, we can integrate all the physiological processes that require carbon (e.g., growth, defense, osmoregulation, reproduction) and provide a more comprehensive cost-benefit assessment.

While our novel results highlight a critical role of stored NSC on plant water relations and a potentially delicate balance between costs and benefits of EM networks under carbon stress, much research is still needed. For instance, we need to elucidate the specific mechanisms (including the involved organic solutes) by which stored NSC depletion decreases osmoregulation, and how these mechanisms vary across species and climate gradients. We also need a better understanding of the potential effect of plant-fungal interactions on NSC depletion and plant water relations in complex EM networks under climate change. For instance, does individual-level (as opposed to stand level) mortality caused by low intensity fire, drought or insects cause rippling effects of similar magnitude to the ones we document when multiple trees are interconnected? We hope that our research motivates such work.

## Methods

### Experimental Design

Ponderosa pine is one of the most widely distributed species in North America and has been extensively used as a representative gymnosperm in ecological, physiological, and forestry studies. In the spring of 2016, we planted eighty, one-year-old ponderosa pine seedlings (source: Zone IV-V block of the Missoula Ponderosa Pine Seed Orchard, Department of Natural Resources) in forty 19 L pots at the University of Montana greenhouse. Each pot contained two seedlings. We used a soil mixture consisting of 40% sand, 30% topsoil, and 30% peat moss, which was kept at field capacity throughout the duration of the experiment to avoid water deficit. Seedlings came already inoculated with ectomycorrhizal Pezizales fungi from source. However, we also inoculated the rhizosphere of each seedling with a mixture of *Rhizopogon* spores (Mycorrhizal Applications. Grants Pass, OR) to ensure colonization. Both *Rhizopogon* and Pezizales form ectomycorrhizal structures that potentially redistribute resources among plants ^20,46^. Half of the pots contained a stainless-steel mesh barrier that separated seedlings (139.7 μm wire diameter and 177.8 μm pore diameter) which was penetrable by hyphae but not roots. The remaining pots contained a Plexiglass barrier to prevent both root and ectomycorrhizal connections between seedlings to create a control treatment and observe patterns in the absence of ectomycorrhizal networks. Both barrier types were secured using silicone sealant. After inoculation, seedlings grew unperturbed for 47 weeks and established mature ectomycorrhizal associations (Figure S1 and S2). Despite our efforts, roots in pots with plexiglass barriers pierced the silicon sealant to form ectomycorrhizal networks. While this precluded a treatment without EM networks, our main goal was to assess how NSC depletion of neighbor seedlings influenced NSC pools and water relations in non-stressed, connected hosts. Because all key variables measured did not differ between pots with and without root connections (Table S1), the two barrier treatments were pooled for statistical analyses. All subsequent treatments described below were applied uniformly in each of the two underground barrier treatments.

In the fall of 2017, we split pots from each treatment into two groups (NSC-depleted and non-depleted) and applied a 3-week NSC depletion treatment to one group using light-blocking covers that reduced photosynthetically photon flux density below compensation point (Fig. 1, Figure S3). In each pot within the NSC-depleted treatment, we placed a cover over one of the two seedlings (dark, D) to inhibit photosynthesis while the other seedling (light paired with dark, LD) was left undisturbed. Each cover consisted of a wire scaffolding (20 x 40 cm) overlaid with aluminum foil that blocked incoming light while keeping air temperatures similar to those in non-covered neighbor seedlings (ca. 24 °C). We pierced 5 mm diameter holes evenly across the cover walls which facilitated air flow, kept the atmosphere around the plant unsaturated at less than 50%, and allowed canopy transpiration. Relative humidity was measured using a Vaisala HMP35C sensor (Vantaa, Finland). We maximized ventilation inside the covers while maintaining PAR below 0.40 μmol quanta m^−2^ s^−1^ which is far below compensation point (Figure S4). We measured PAR with a Licor LI-190SA Quantum sensor (Lincoln, NE, USA). Combining shading with ectomycorrhizal networks allowed us to alter the overall carbon budget of a networked system and indirectly modify NSC pools in non-shaded plants. This procedure generated non-depleted networks with both connected seedlings in the light (light, LL) and NSC-depleted networks with one seedling in the light (LD) connected to one seedling in the dark (D). After three weeks of light deprivation, we randomly selected plants from ten pots (five from each underground barrier treatment) in both NSC-depleted and non-depleted treatments to assess physiological status (see below) of LL, LD, and D seedlings and harvested them to establish base-line ^13^C/^12^C ratios (*δ*^13^C) in all organs across seedling types. We then labeled with ^13^C (see below) the remaining LD seedlings from NSC-depleted networks and one of the two LL seedlings (chosen at random) from the non-depleted NSC networks to assess potential carbon transfer between individuals in the presence and absence, respectively, of carbon-limiting conditions seven days later. Comparing carbon transfer between non-depleted and NSC-depleted networks let us assess how carbon transfer responds to carbon limitation. One week after the labeling, we re-assessed physiological status and harvested the remaining seedlings in both treatments to measure changes in *δ*^13^C across all organs (Fig. 1).

**Fig. 1.**
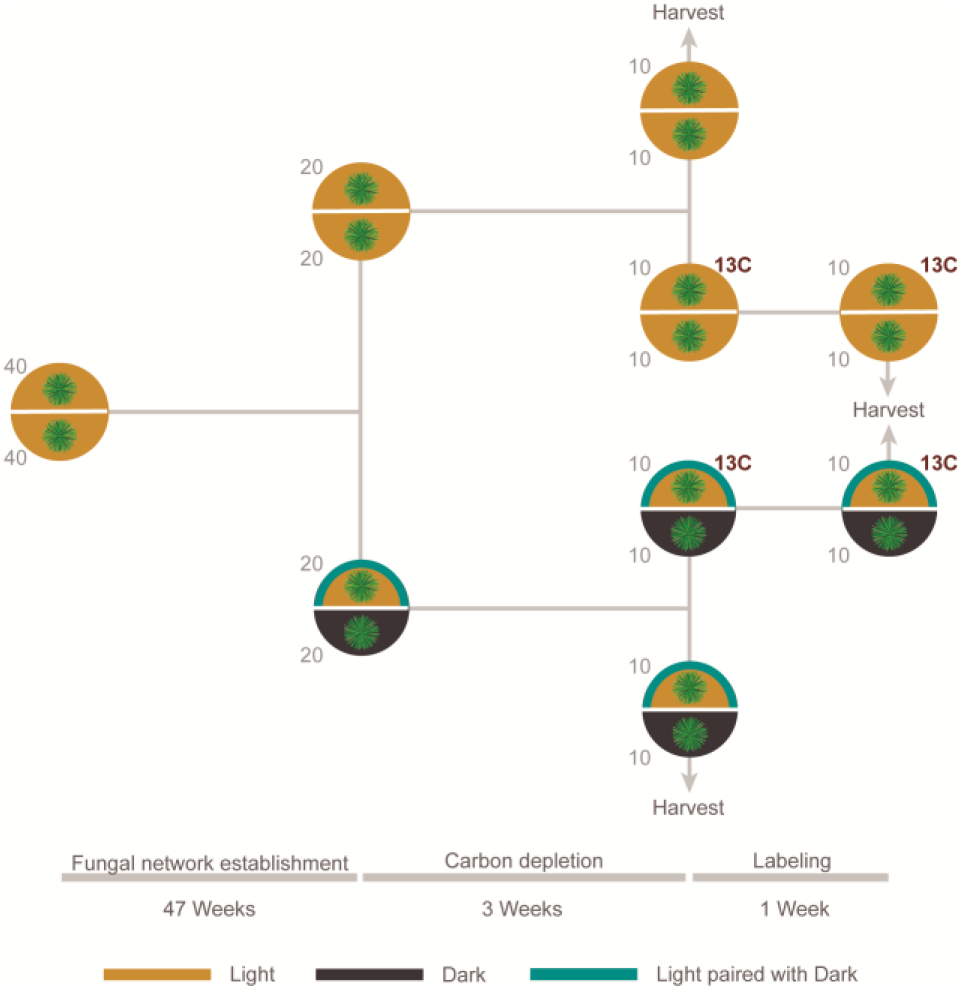
Experimental design. Circles represent pots with barriers and a seedling on each side. Colors represent LL seedlings exposed to natural light (golden), D seedlings with light-blocking covers (black), and LD seedlings exposed to natural light paired with plants with covers (teal border). Numbers on the side of each circle indicate the sample size of the seedling type represented in that side of the pot at a given point on time. Red 13C annotations indicate pots labeled with ^13^C. Arrows indicate harvesting events.

**Fig. 2.**
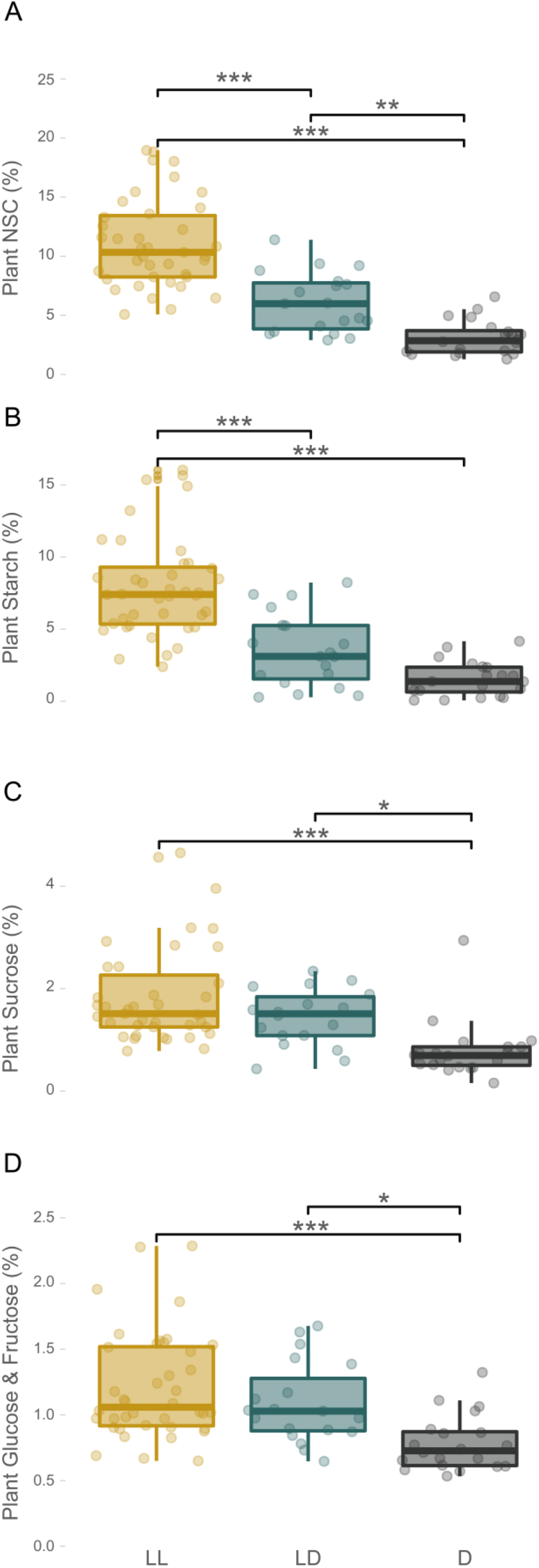
Stored NSC decreased in seedlings of the NSC-depletion treatment. Panels correspond to whole-plant concentrations in percent dry weight. Colors represent LL seedlings exposed to natural light (golden), D seedlings with light-blocking covers (black), and LD seedlings exposed to natural light paired with plants with covers (teal). Lines within boxes represent the median and top and bottom hinges represent 25th and 75th percentiles. Whiskers indicate highest and lowest value no further than 1.5 times the inter-quartile range represented by the hinges. Dots represent the distribution of the data. Asterisks indicate the degree of significance between groups (* = 0.05, ** = 0.01, *** = < 0.001).

**Fig. 3.**
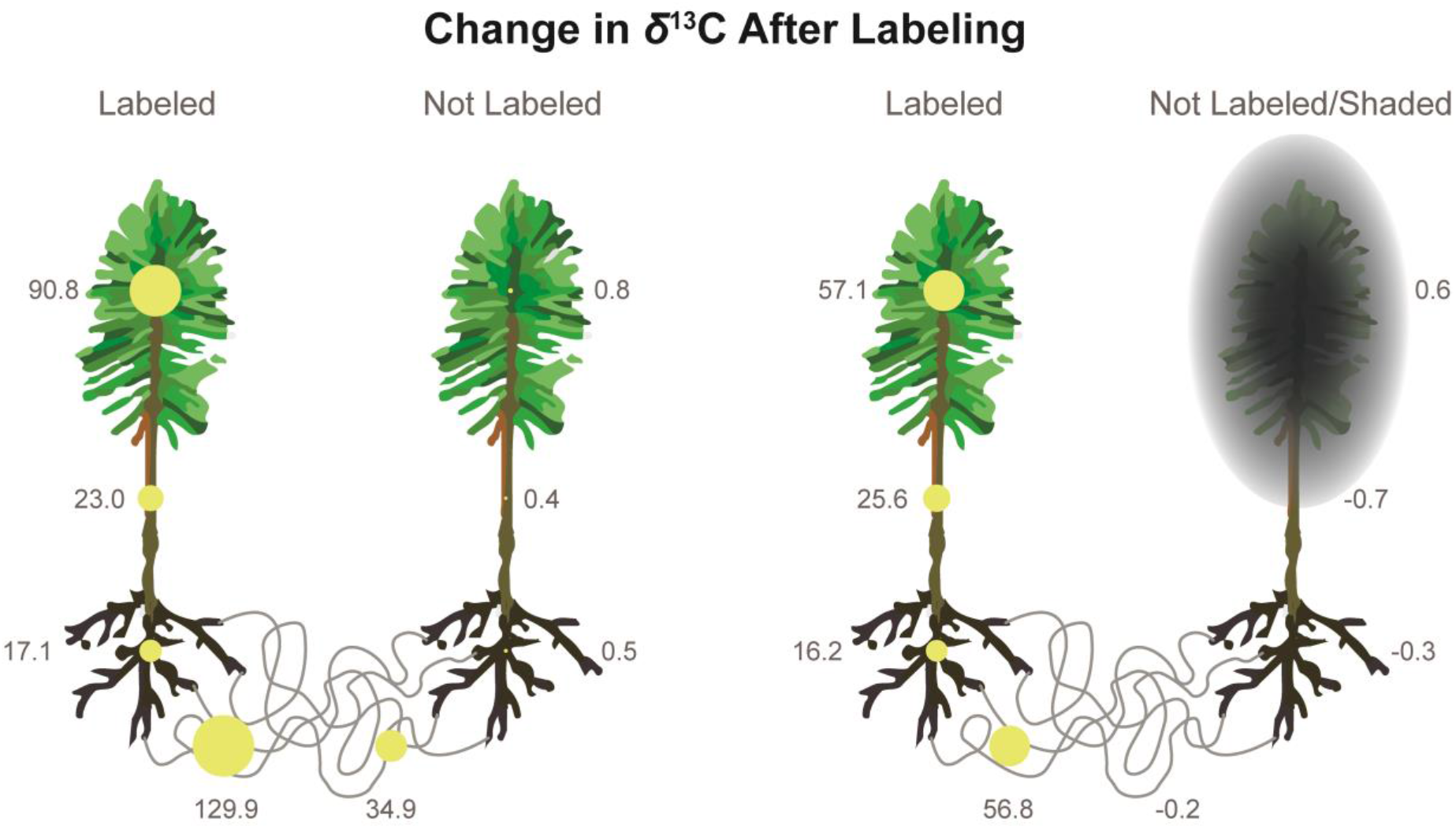
^13^C isotope reached all organs in labeled plants and the ectomycorrhizal network but was not transferred to NSC-depleted hosts. The area of the yellow circles represents the average change in ^13^C/^12^C ratios in each tissue as a result from labeling (mean tissue *δ*^13^C after labeling – mean tissue *δ*^13^C before labeling) in parts per thousand (‰; also, annotated besides each circle). Positive values indicate an increase in *δ*^13^C. See Figure S7 and S8 for raw *δ*^13^C values for each tissue and labeling time, confidence intervals, and statistical significances.

**Fig. 4.**
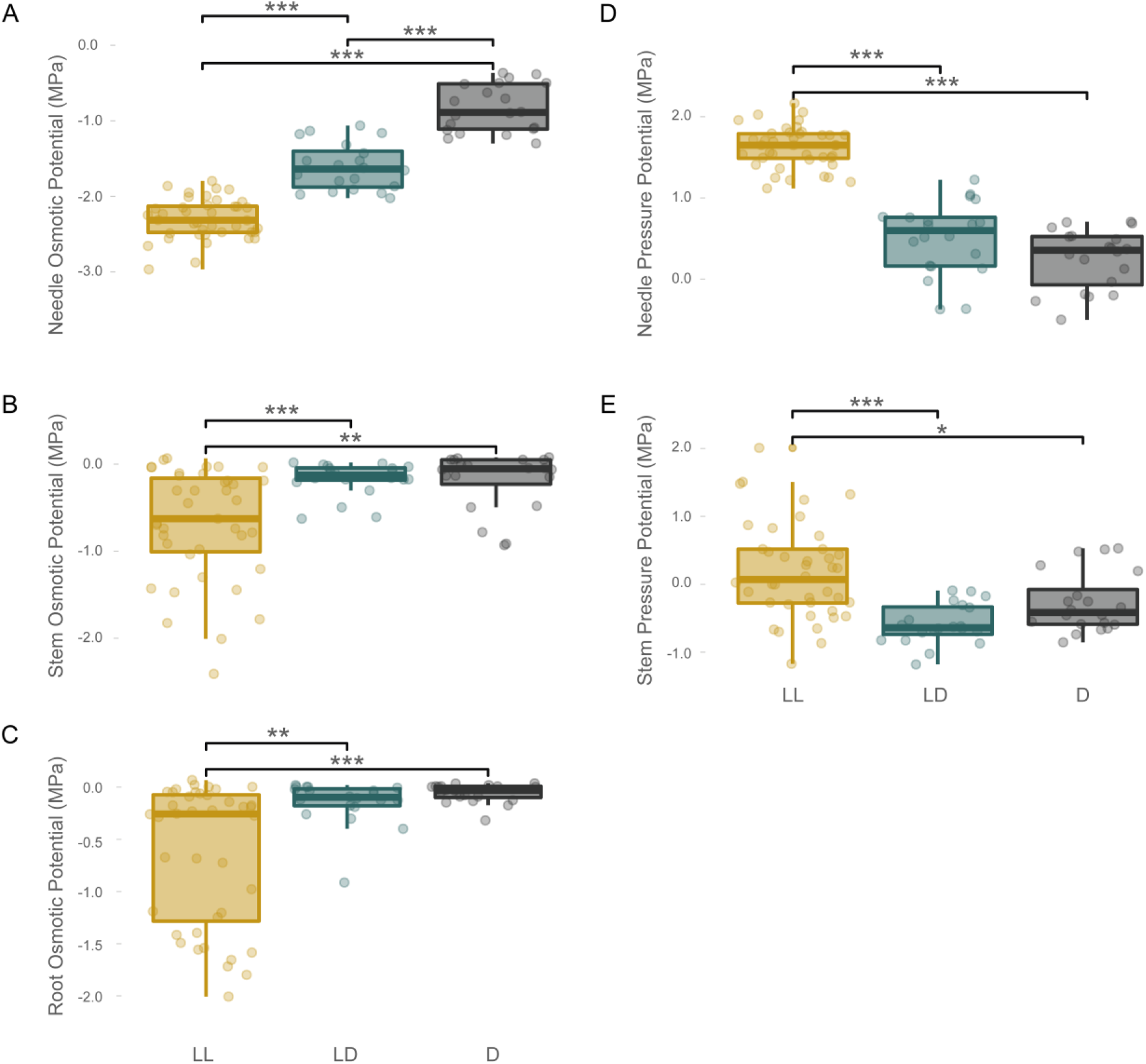
All organs in NSC-depleted seedlings exhibited an increase of osmotic potential and a decrease in turgor. Panels on the left correspond to osmotic potentials of A) needles, B) stems, and C) roots. Panels on the right correspond to pressure potentials of D) needles and E) stems. Colors represent LL seedlings exposed to natural light (golden), D seedlings with light-blocking covers (black), and LD seedlings exposed to natural light paired with plants with covers (teal). Lines within boxes represent the median and top and bottom hinges represent 25th and 75th percentiles. Whiskers indicate highest and lowest value no further than 1.5 times the inter-quartile range represented by the hinges. Dots represent the distribution of the data. Asterisks indicate the degree of significance between groups (* = 0.05, ** = 0.01, *** = < 0.001).

**Fig. 5.**
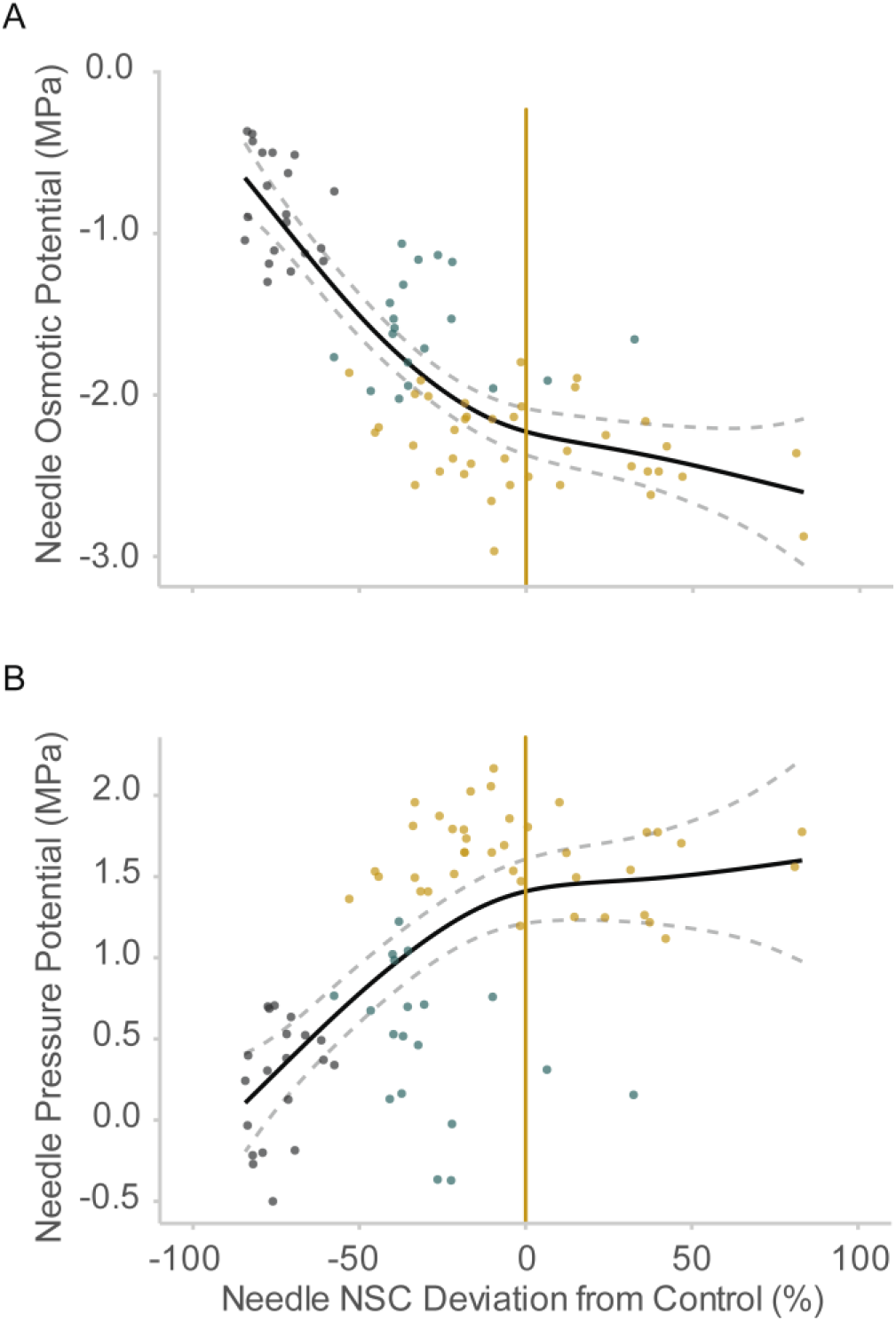
Water relations are impaired when NSC drop below values corresponding to non-stressed plants. Panels correspond to A) osmotic potential and B) turgor pressure. All seedling types are merged for this analysis. Vertical golden lines indicate average leaf NSC content in non-depleted pots (i.e., light plants). A loess function was fit to the data to best represent the relationship between variables. Dashed lines indicate 95% confidence interval of the regression lines.

**Fig. 6.**
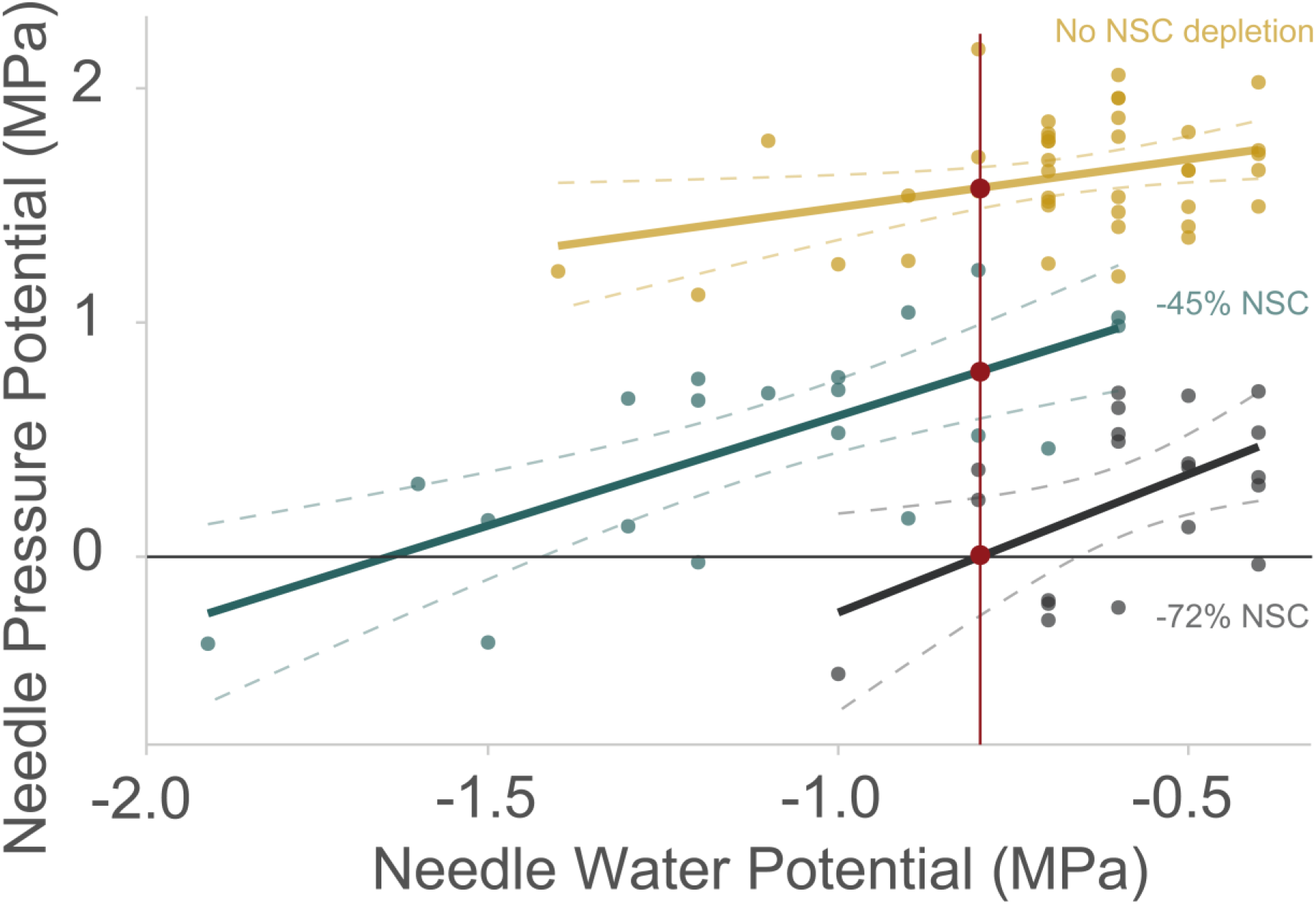
NSC depletion is associated with turgor loss at high water potentials. Colors represent LL seedlings exposed to natural light (golden), D seedlings with light-blocking covers (black), and LD seedlings exposed to natural light paired with plants with covers (teal). Vertical red line and points indicate differences in turgor at a given water potential. Horizontal line indicates turgor loss point. Dashed lines indicate 95% confidence interval of the regression lines.

### Isotopic Labeling and Carbon Transfer Variables

Labeling was performed following similar methods to Song *et al.* (2015). We enclosed the entire canopy of each labeled plant in a clear 2 L plastic bag. Each bag was injected with 20 mL of Carbon-^13^C dioxide (99 atom % 13C, <3 atom % 18O, Sigma Aldrich, (St. Louis, MO, USA)) for a ratio of 10 mL of ^13^C·L^−1^ of air and was left undisturbed for 2 hours. ^13^C injections were performed at midday over two consecutive days. After each labeling event, enriched air was removed from bags and directed outside the greenhouse facilities using an industrial vacuum, thus preventing contamination of neighboring seedlings. Additionally, extra seedlings were randomly interspersed throughout the treatments and their needles were used to assess possible contamination during labeling. No signs of increased *δ*^13^C were observed in these seedlings. Ground samples of dry needles, stems, and roots of all remaining seedlings (labeled or not) were sent to the Stable Isotope Facility at University of California, Davis and analyzed to obtain *δ*^13^C values. We also collected a representative amount of fungal material from ectomycorrhizal root tips of LL, LD, and D seedlings (5 samples) before the labeling event to obtain base-line *δ*^13^C fungal values as well as several labelled LL (4) and LD (3) seedlings, and non-labelled LL (9), and D (10) seedlings after labeling and analyzed their *δ*^13^C. Fungal samples were collected to ensure that the applied ^13^C label traveled from needles to fungal tissue on the labeled side as well as to fungal tissue of the paired, non-labeled plants connected through the ectomycorrhizal network. These descriptive measurements served as an extra validation of the existence of an ectomycorrhizal network between seedling pairs already observed at the time of harvest (Figure S1).

### Measurements of water relations

After re-acclimating D seedlings to light for 2 hours (see Figure S5, panel F), we measured assimilation, stomatal conductance, and respiration in each seedling with a Licor 6400XT (Lincoln, NE, USA). We then measured midday leaf and stem water potential (WP) (i.e., xylem tension due to water deficit) immediately before harvesting (Fig. 1) using a pressure chamber (PMS Instrument Company, (Corvallis, OR, USA)) following methods in Kaufmann (1968). Stem WP was estimated by equilibrating the WP of a needle bundle with that of the stem following methods from Begg & Turner (1970) and measuring the equilibrated bundle as above. Subsequently, we harvested seedlings and collected tissue samples (needles, stem, and roots) for osmotic potential measurements (i.e., water retention capacity via solutes potentially derived from NSC). Tissue samples were wrapped in aluminum foil to prevent artificial changes in osmotic potential due to water loss or photosynthetic activity, placed in small Ziploc bags, frozen in dry ice and transported to the lab within 2 hours. The rest of the seedling was placed in a Ziploc bag containing a wet paper towel partially covered in tin foil to prevent desiccation without introducing external moisture into the tissues in contact with the towel ^32^, placed in a cooler, and transported to the lab. There, we measured relative water content (RWC) and hydraulic conductivity (i.e., capacity to transport water ^49^). We describe the methods used to measure water content and hydraulic conductivity in Methods S1. We also used the rest of the seedling to measure plant NSC pools (see below). Upon arrival to the lab, samples collected for osmotic potential were pressed to extract cellular liquid contents and the expressed solution was used to saturate 28 mm^2^ filter paper disks following methods from Grange (1983) ^50^. This method avoids potential osmotic artifacts due to starch mobilization or low sample water content. Disks were then placed in a C-52 sample chamber attached to a Psypro data logger (Wescor, Inc. Logan, UT, USA) to measure osmotic potential. Finally, pressure potential (i.e., positive pressure exerted by cells against cell walls when turgid ^12^) was calculated in stems and needles as the difference between their respective water and osmotic potential. Pressure potential was not calculated in roots because we lacked root WP measurements.

### Non-structural carbohydrates

Non-structural carbohydrates were analyzed in all tissue types (needles, stems, and roots) collected at harvest. A sample of each tissue was microwaved for 180 seconds at 900 Watts in three cycles of 60 seconds to stop metabolic consumption of NSC pools, and oven-dried at 70 °C. Samples were dried to a constant mass and finely ground into a homogenous powder. Approximately 11 mg of needle tissue and 13 mg of stem or root tissue were used to analyze NSC dry mass content following the procedures and enzymatic digestion method from Hoch, Popp & Körner (2002) and Galiano, Martínez-Vilalta, Sabaté & Lloret (2012). NSC, starch, sucrose, and glucose + fructose tissue concentrations were then multiplied by their respective tissue fraction (the dry mass of a tissue relative to whole-plant dry mass) to obtain whole-plant concentrations. Whole-plant dry mass for each seedling was calculated by combining the dry mass of all samples and the remaining biomass. These methods are described in Methods S1.

### Statistical Analyses

We observed no differences between pots with (Plexiglas barrier) and without (mesh barrier) root connections for all key variables measured (Table S1), and data from the two barrier treatments were pooled. We tested differences among seedling types (LL, LD, D) in all variables using ANOVA and Tukey’s Honestly Significant Difference tests. Note that sample size of LL seedlings is twice that of LD and D seedlings because both plants in non-depleted networks lack of covers and are LL seedlings (Fig. 1). However, because treatment comparisons excluding a random half of the LL seedlings yielded the same results, we decided not to exclude any data. To test differences in NSC depletion, we compared total NSC concentration and concentrations of each NSC component (i.e., starch, sucrose, and glucose and fructose together) among seedling types at both organ and whole-plant levels. To test the effectiveness of the labeling process and assess carbon translocation among organs, for each organ and treatment, we compared *δ*^13^C values after labelling to those prior to labeling. Note that D seedlings were excluded because they were never labeled. We only analyzed a few fungal samples for *δ*^13^C (see above) as an additional check that the ^13^C label could travel from the labelled seedling to the fungal network of the non-labelled seedling, thus ensuring that plants were indeed connected via fungal networks. Likewise, to assess potential carbon transfer to non-labeled seedlings, for each organ and treatment, we compared *δ*^13^C values after labeling the paired seedling to those before labelling. LD seedlings were excluded because they were all labeled. We also tested differences in hydraulic conductivity, WP, osmotic potential, pressure potential and relative water content in each organ for which we collected data among seedling types.

To assess both direct and indirect effects of NSC depletion on water relations, we assessed whether osmotic potential and pressure potential changed in response to NSC depletion. We used two linear regressions with needle % NSC deviation from control as predictor and either leaf osmotic or pressure potential as response variables. We focused on needles because they are the organ with most living cells exposed to dry conditions and, therefore, most susceptible to turgor loss. Finally, we assessed whether NSC depletion influences turgor loss as WP decreases using linear regressions. Leaf pressure potential was the response variable and the interaction between leaf WP and whole-plant % NSC deviation from control as predictor. In this analysis, we focus only on differences in intercepts among NSC levels (i.e. NSC effect on leaf WP at turgor loss point, a broadly used indicator of drought tolerance) and among their slopes (i.e. rate of turgor loss per unit change in water potential) because a significant relationship between leaf pressure potential and leaf WP is expected given that the former is calculated from the later.

## Acknowledgements

This work was partially supported by a National Science Foundation grant to AS (BCS 1461576). GS received funding from the NSF Experimental Program to Stimulate Competitive Research (EPSCoR) Track-1 EPS-1101342 (INSTEP 3). PD received funding from NSF EPSCoR RII Track 1 award number IIA-1443108. YL is grateful to MPG Ranch for funding. The authors thank Laura Thornton and Maria Pilar de Moreta for their help collecting data, and Mauri Valett and Lila Fishman for letting us use their facilities. The authors also thank D. Stanton, R. Montgomery, R. Koide, and D. Ulrich for their comments in early versions of this manuscript.

