## Supplementary material for "Plant carbohydrate depletion impairs water relations and spreads via ectomycorrhizal networks": Figure S

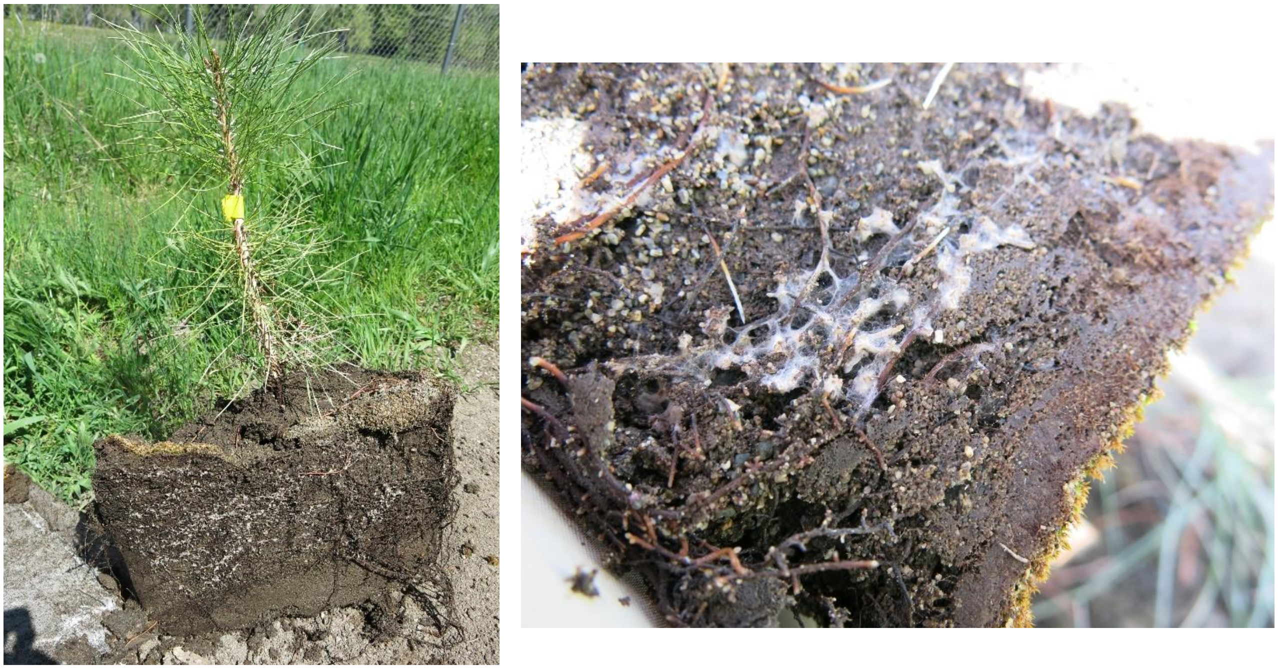

**Fig. S1.** Two examples of the typical underground profile observed in pots with mesh barriers at harvesting. The flat surface observed here corresponds to the area in contact with the mesh barrier. Notice how hyphae are highly abundant at the area in contact with the mesh. This pattern was always mirrored in the other side of the mesh indicating that hyphae from both seedlings were at least in contact and most likely the same.

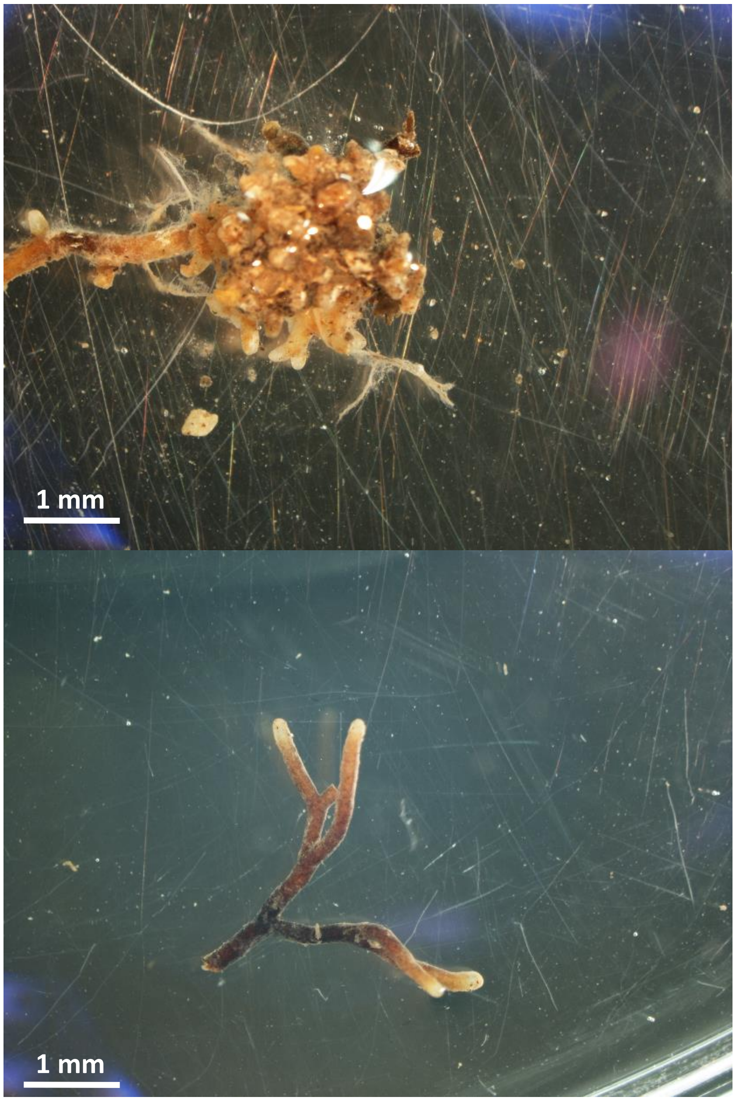


**Fig. S2.** Root tips colonized by ectomycorrhizae. Notice the bifurcated morphology of the root tips typical from roots infected by mycorrhizae.

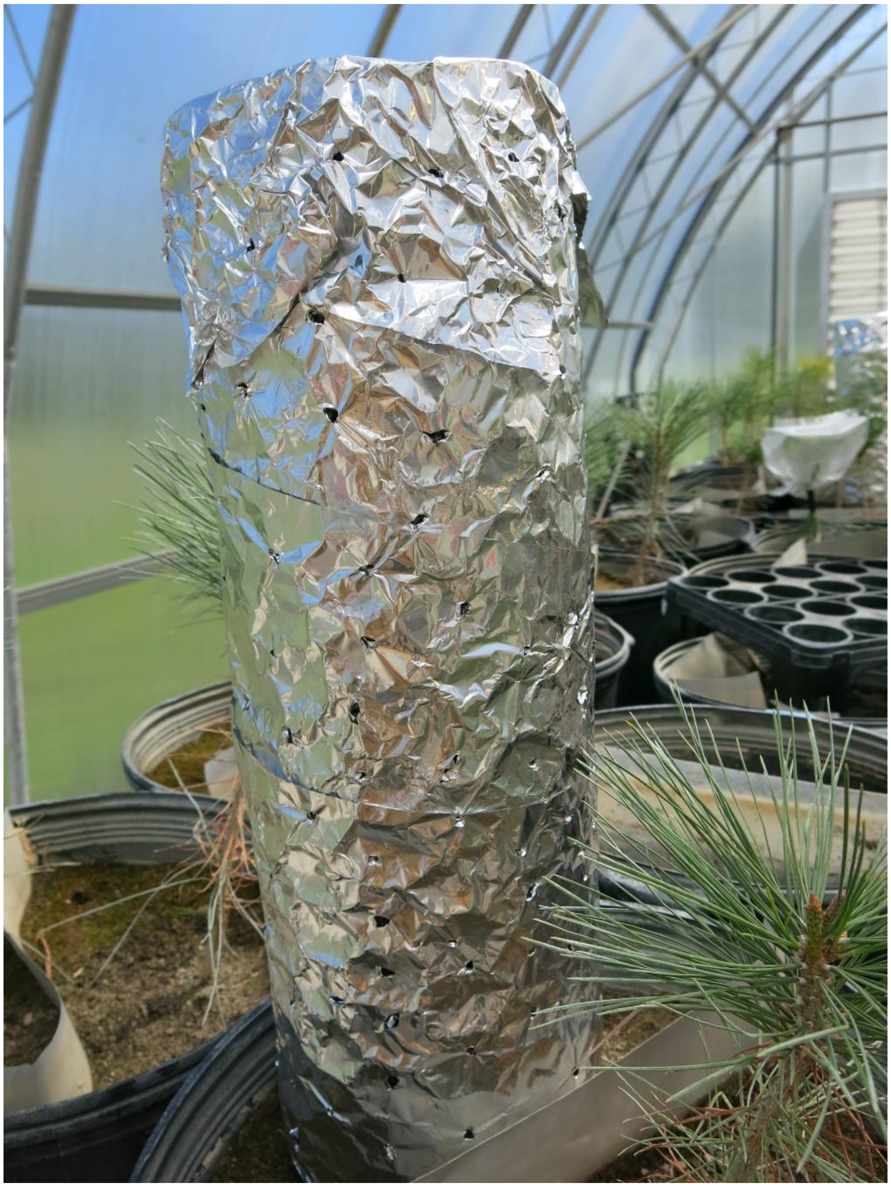

**Fig. S3.** Detail of the light-blocking covers used to impose non-structural carbohydrate depletion.

**Table S1.** No differences between pots with and without root connections allow merging of both groups. Based on Student's t-test. Variables were properly transformed when needed to meet test assumptions.

| Variable compared between pots with and without root connections | P-value | Explanation of differences |
|---|---|---|
| Canopy biomass | 0.868 | |
| Root biomass | 0.013 * | More roots in root barrier to pierce it and reach other side of pot to create network |
| Assimilation | 0.327 | |
| Stomatal Conductance | 0.902 | |
| Respiration | 0.016 * | Higher in root barrier likely because more roots |
| Plant NSC Deviation from Control | 0.848 | |
| Plant Total NSC Concentration | 0.440 | |
| Plant Starch Concentration | 0.486 | |
| Plant Sucrose Concentration | 0.469 | |
| Plant Glucose + Fructose Concentration | 0.950 | |
| Leaf Relative Water Content | 0.853 | |
| Leaf Water Potential | 0.060 | |
| Leaf Osmotic Potential | 0.336 | |
| Leaf Pressure Potential | 0.778 | |
| Leaf Saturated Water Content | 0.603 | |
| Leaf Osmotic Potential at Full Turgor | 0.845 | |
| Leaf Water Potential at Turgor Loss | 0.837 | |
| Leaf Relative Water Content at Turgor Loss | 0.908 | |
| Leaf Capacitance at Full Turgor | 0.245 | |
| Leaf Capacitance at Turgor Loss | 0.060 | |
| Leaf Modulus of Elasticity | 0.538 | |
| Leaf NSC Deviation from Control | 0.123 | |
| Leaf Total NSC Concentration | 0.123 | |
| Leaf Starch Concentration | 0.109 | |
| Leaf Sucrose Concentration | 0.366 | |
| Leaf Glucose + Fructose Concentration | 0.348 | |
| Stem Relative Water Content | 0.448 | |
| Stem Water Potential | 0.310 | |
| Stem Osmotic Potential | 0.728 | |
| Stem Pressure Potential | 0.481 | |
| Stem hydraulic Conductivity | 0.412 | |
| Stem Total NSC Concentration | 0.521 | |
| Stem Starch Concentration | 0.231 | |
| Stem Sucrose Concentration | 0.642 | |
| Stem Glucose + Fructose Concentration | 0.686 | |
| Root Relative Water Content | 0.966 | |
| Root Osmotic Potential | 0.389 | |
| Root hydraulic Conductance | 0.132 | |
| Root Total NSC Concentration | 0.599 | |
| Root Starch Concentration | 0.341 | |
| Root Sucrose Concentration | 0.932 | |
| Root Glucose + Fructose Concentration | 0.712 | |

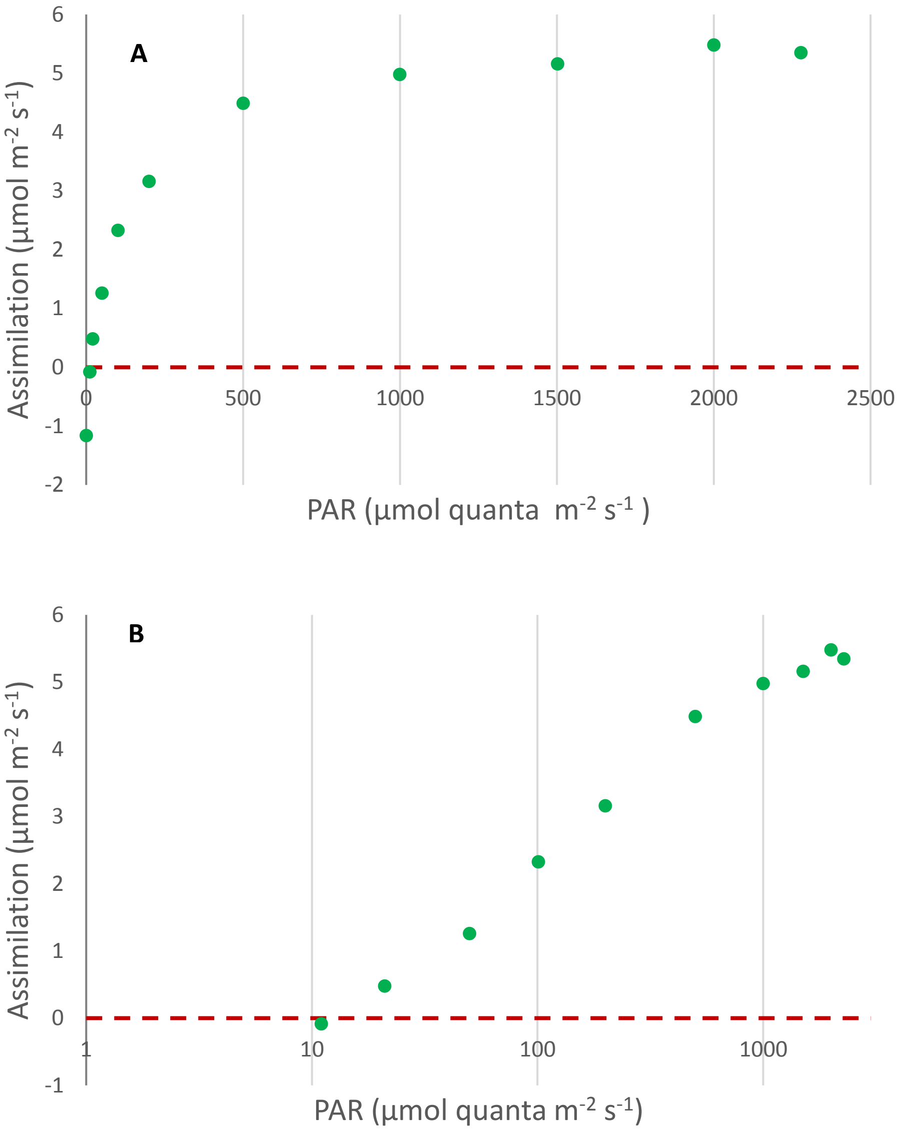


**Fig. S4.** Light curve showing the amount of Photosynthetic Active Radiation (PAR) necessary to reach compensation point (dashed red line). Panel A shows the curve with natural axis values. Panel B shows PAR in a log10 axis to better visualize values near compensation point. Compensation point occurred at ca. 11 µmol quanta $m^{-2}$ $s^{-1}$. Light-blocking covers reduced PAR to 0.40 µmol quanta $m^{-2}$ $s^{-1}$. Thus, ensuring that they would lead to depletion of stored non-structural carbohydrates.

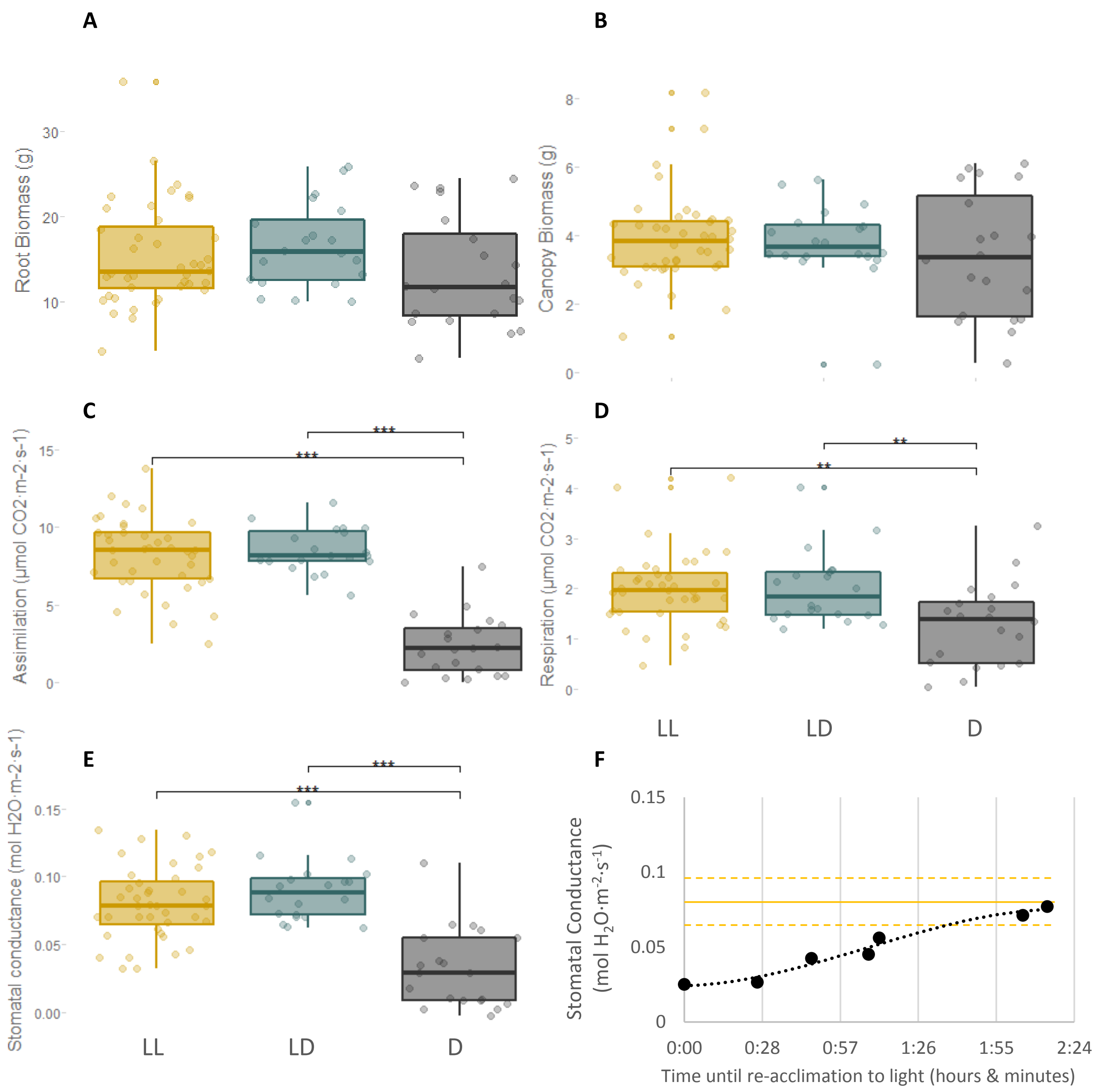


**Fig. S5.** NSC depletion in LD seedlings is most likely explained by an increase in fungal NSC demand. There was no significant growth, increase in respiration, or reduction in assimilation relative to LL seedlings that could alternatively explain NSC depletion. D seedlings were the only treatment to show reduced respiration (likely to avoid total depletion of NSC pools) and reduced assimilation (likely due to lower stomatal conductance in response to low turgor). We ensured that gas exchange measurements in D seedlings were not affected by recent exposure to dark by re-acclimating D seedlings to light conditions before measurements (see panel F and its explanation in caption). Panel A and B corresponds to below- and above-ground biomass. respectively. Panel C to E correspond to leaf net assimilation, respiration, and stomatal conductance, respectively. Panel F shows that 2 hours is enough to bring stomatal conductance of a representative extra seedling exposed to dark for 24 hours within the $25^{th}$ -$75^{th}$ percentile range (dashed golden lines) and close to the mean (solid golden line) of a light-acclimated plant, thus eliminating any artificial effects of darkness on gas exchange measurements. Hence, any differences in gas exchange after this re-acclimation time are due to treatment effects. Colors represent LL seedlings exposed to natural light (golden), D seedlings with light-blocking covers (black), and LD seedlings exposed to natural light paired with plants with covers (teal). Lines within boxes represent the median and top and bottom hinges represent 25th and 75th percentiles. Whiskers indicate highest and lowest value no further than 1.5 times the inter-quartile range represented by the hinges. Dots represent the distribution of the data. Asterisks indicate the degree of significance between groups (* = 0.05, ** = 0.01, *** = < 0.001).

**Methods S1.** Description of methods used for relative water content, hydraulic conductivity, and non-structural carbohydrates (NSC).

Relative water content

We used a sample of roots, stems, and needles of each seedling to measure their relative water content (RWC). First, samples were weighted to obtain fresh weight and returned to Ziploc bags in the cooler to avoid changes in hydraulic conductivity due to exposure to dry air. For consistency, root fresh weight was measured before any other tissue to avoid changes in RWC or hydraulic conductivity due to exposure to dry air. After hydraulic conductivity measurements (see below), stem, needle, and root samples were hydrated to full turgidity for 5 hours in a water bath at 10 °C. After hydration, we blotted each sample to remove surface moisture using a paper towel and weighed them to determine weight at full turgor. Samples were then oven dried at 70ºC, until a constant mass was achieved and weighed to determine dry weight. RWC was calculated as: ((Fresh weight - Dry weight)/(Turgid weight - Dry weight))*100 following methods from [1]. The rest of the seedling was dried, separated by organ, and weighed. These weights were later combined with sample dry weights to calculate whole plant RWC by multiplying the dry mass of each tissue relative to whole-plant dry mass (i.e., tissue fraction) by their respective RWC. Whole-plant dry mass for each seedling was calculated by combining the dry mass of all samples and the remaining biomass.

Hydraulic conductivity

We measured stem hydraulic conductivity and root hydraulic conductance using the gravimetric method [2], after fresh weight measurements. We used the same hydraulic apparatus described in Sapes (2019)[3] capable of measuring hydraulic conductance of both whole root systems and stems. After measuring fresh weight, stem segments were immersed in deionized water for 20 minutes to relax xylem tensions that could artificially alter conductivity values [4]. After relaxation, stems were relocated to the hydraulic apparatus and each end was recut twice at a distance of 1 mm from the tips each time (total of 2 mm per side) to remove any potential emboli resulting from transport, previous cuts, and relocation [5]. Stems were then connected to the hydraulic apparatus while under water, with their terminal ends facing downstream flow. The stems were then raised out of the water and the connections were checked to ensure that there were no leaks. A solution of water with 10 mM KCl degassed at 3 kPa for at least 8 hours was then used for all hydraulic measurements [6]. First, initial background flow was measured to account for the flow existing under no pressure, which can vary depending on the degree of dryness of the measured tissue [7–9]. Second, a pressure gradient of 5-8 kPa was applied to run water through the stem and pressurized flow was measured. This small pressure gradient prevented embolism removal from the samples while ensuring flow. Lastly, final background flow was measured, initial and final background flows were averaged, and flow was calculated as the difference between pressurized flow and average background flow. Native specific hydraulic conductivity (K) was estimated in stems as the flow divided by the pressure gradient used and standardized by xylem area and length. Stem segments where then removed from the apparatus and placed in a water bath for measurements of RWC (see above). The configuration of the apparatus was then changed to measure whole root system hydraulic conductance using the same gravimetric principle as explained in Sapes (2019)[3]. Flow, including initial and final background flow, was measured as above and whole root native hydraulic conductance (k) was estimated as the flow divided by the pressure gradient used and standardized by xylem area at the root collar. We used the R code published in Sapes (2019)[3], see Methods S1 in Supporting Information) to calculate pressurized and background flows once flow stabilizes. Once flow rates were measured, root samples were placed in a water bath to be used in measurements of RWC.

Non-structural carbohydrates

Sample powder was dissolved in 1.6 mL of distilled water and incubated at 100 °C for 60 min to extract carbohydrates. An aliquot of the extract was used to determine soluble sugar concentrations (i.e., glucose, fructose, and sucrose) through enzymatic conversion of sucrose and fructose into glucose by invertase from *Saccharomyces cerevisiae* and phosphoglucose isomerase, respectively (I4504 and G3293, Sigma-Aldrich). Total *NSC* concentration was obtained from another aliquot incubated in amyloglucosidase from *Aspergillus niger* (10115, Sigma-Aldrich) at 50 °C during 16 hours to break down all *NSC* (starch included) into glucose. In both cases, the concentration of glucose was determined photometrically in a 96-well microplate reader (BioTek™ EL800, Winooski, United States) after enzymatic conversion of glucose into glucose-6-phosphate by glucose hexokinase. The dehydrogenation of glucose causes an increase in optical density at 340 nm. All the NSC and soluble sugar concentrations were expressed as percentages of dry matter. Although the quantification of NSC has been proven difficult and inconsistent among laboratories, the reasonable consistency within a given laboratory allows comparisons among samples [10]. We calculated the total pool of NSCs, starch, soluble sugars, and glucose or fructose in each tissue by multiplying the corresponding concentration of each tissue by its dry weight. Concentrations (total NSC and each individual component) were later scaled up to whole-plant level as explained in the RWC section. Finally, the relative degree of NSC depletion of each individual (here on percent NSC deviation from control) was calculated at both tissue and whole-plant levels by subtracting the mean NSC concentration of LL seedlings (i.e. controls) from each individual value and dividing that difference by the mean. This ratio was then multiplied by 100 to represent it as a percentage. Percent NSC deviation from control can range from negative to positive values with negative values expressing NSC depletion and positive values expressing greater NSC pools than the average non-stressed seedling [11].

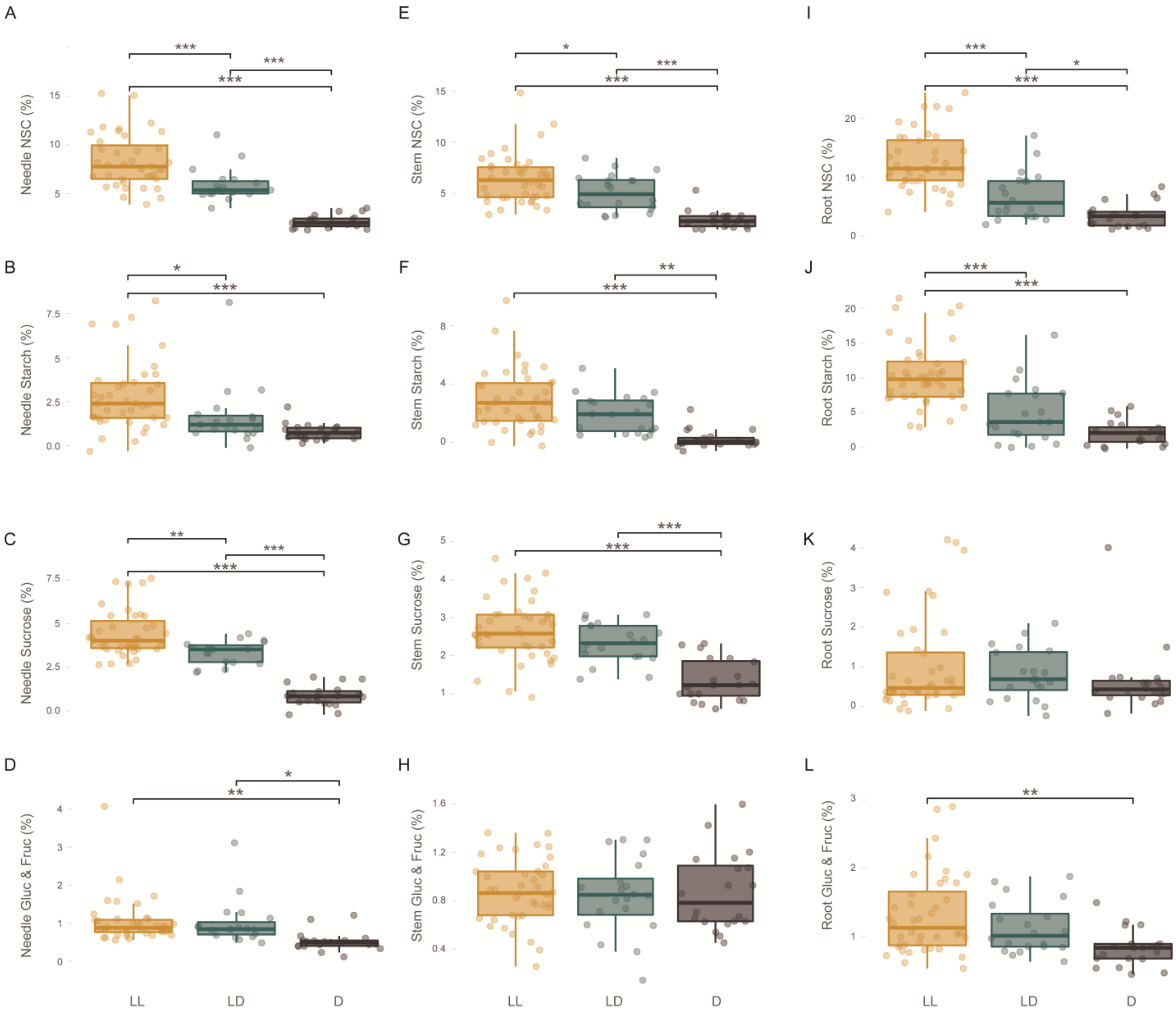
A
B
C
D
E
F
G
H
I
J
K
L
Needle NSC (%)
Needle Starch (%)
Needle Sucrose (%)
Needle Gluc & Fruc (%)
Stem NSC (%)
Stem Starch (%)
Stem Sucrose (%)
Stem Gluc & Fruc (%)
Root NSC (%)
Root Starch (%)
Root Sucrose (%)
Root Gluc & Fruc (%)
LL
LD
D

**Fig. S6.** All seedlings within the NSC-depletion treatment consistently showed low levels of all NSC components across organs. Rows correspond to NSC, starch, sucrose, and Glucose & Fructose concentrations, respectively. Columns correspond to needles, stems, and roots, respectively. Colors represent LL seedlings exposed to natural light (golden), D seedlings with light-blocking covers (black), and LD seedlings exposed to natural light paired with plants with covers (teal). Lines within boxes represent the median and top and bottom hinges represent 25th and 75th percentiles. Whiskers indicate highest and lowest value no further than 1.5 times the inter-quartile range represented by the hinges. Dots represent the distribution of the data. Asterisks indicate the degree of significance between groups (* = 0.05, ** = 0.01, *** = < 0.001).

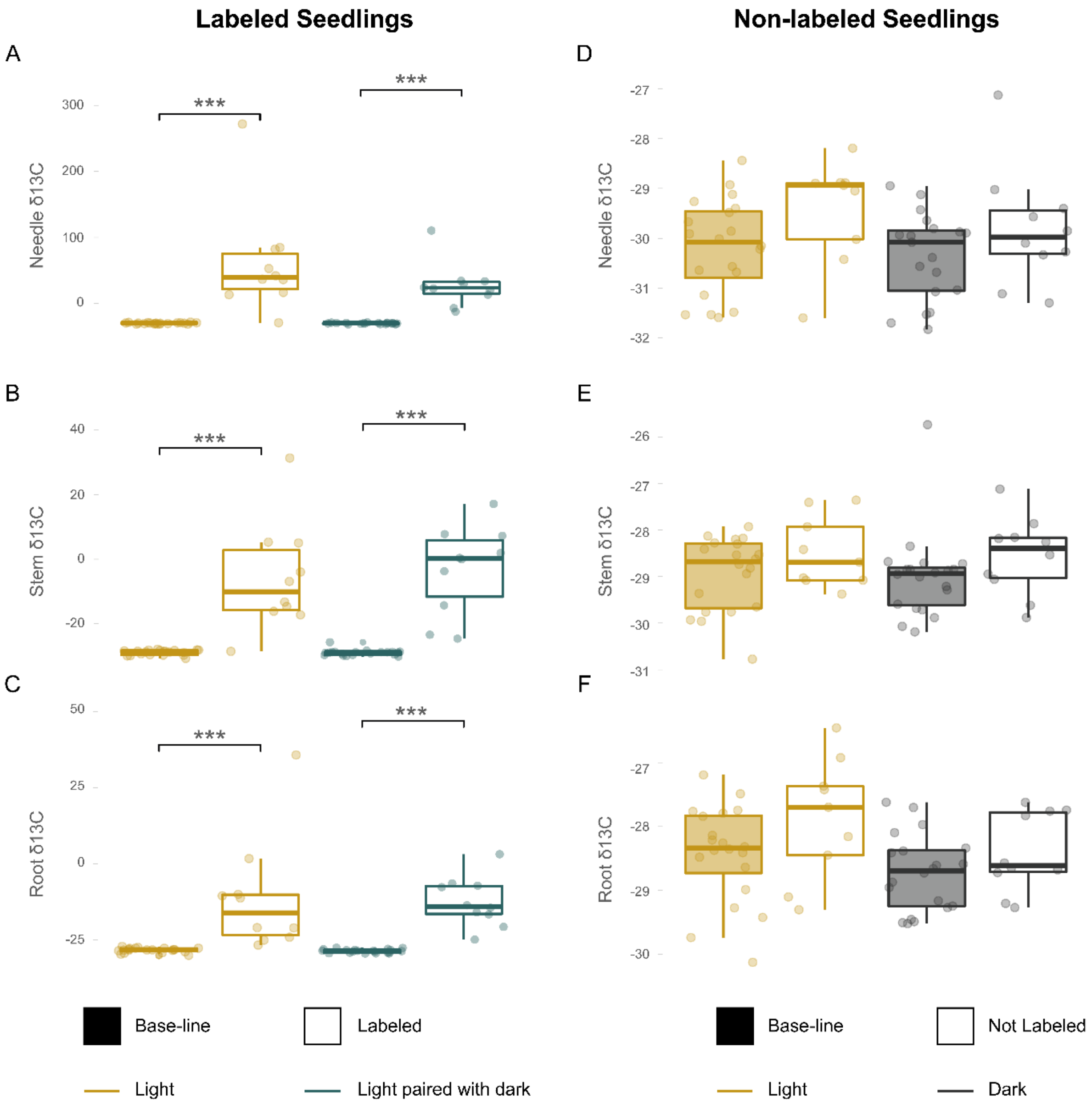


**Fig. S7.** $^{13}C$ isotope reached all organs in labeled plants but was not transferred to carbon-depleted hosts. Panels on the left correspond to $^{13}C/^{12}C$ ratios ($\delta^{13}C$) in A) needles, B) stems, and C) roots, from labeled plants. Panels on the right correspond to $\delta\ ^{13}C$ in D) needles, E) stems, and F) roots, from non-labeled plants. Solid boxes indicate values before labeling (i.e., base-line) and open boxes indicate values after labeling. Colors represent plants exposed to natural light (golden), plants with light-blocking covers (black), and plants exposed to natural light paired with plants with covers (teal). Lines within boxes represent the median and top and bottom hinges represent

25th and 75th percentiles. Whiskers indicate highest and lowest value no further than 1.5 times the inter-quartile range represented by the hinges. Dots represent the distribution of the data. Asterisks indicate the degree of significance between groups (* = 0.05, ** = 0.01, *** = < 0.001). A synthesized version of this figure is provided in Figure 3 of the main manuscript.

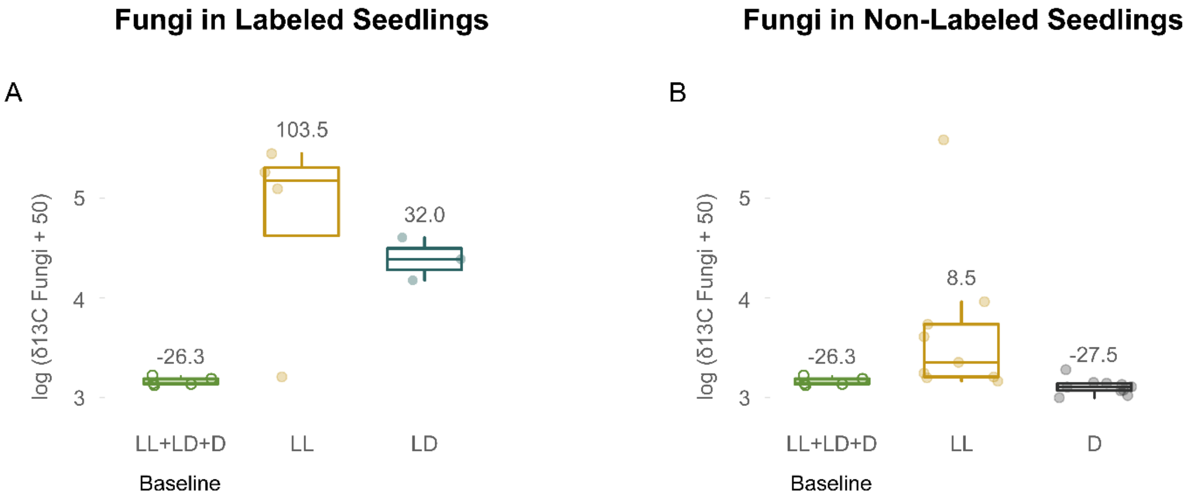


**Fig. S8.** $^{13}C$ isotope only reached both sides of the fungal network in non-NSC-depleted pots as indicated by the enriched values only in LL non-labeled seedlings. Panels on the left correspond to $^{13}C/^{12}C$ ratios ($\delta^{13}C$) Panels correspond to A) fungi associated with labeled seedlings and B) fungi associated with non-labeled seedlings. Notice that a constant has been added to all values in the y-axis to be able to represent negative values in a log scale. For easier interpretation, we provide the actual average values of $\delta\,^{13}C$ for each group above boxplots. Solid green boxes indicate values before labeling pooled across all seedling types (i.e., baseline). Open boxes indicate values after labeling in fungi from plants exposed to natural light (golden), plants with light-blocking covers (black), and plants exposed to natural light paired with plants with covers (teal). Lines within boxes represent the median and top and bottom hinges represent 25th and 75th percentiles. Whiskers indicate highest and lowest value no further than 1.5 times the inter-quartile range represented by the hinges. Dots represent the distribution of the data. Changes in $\delta^{13}C$ in fungal tissue are descriptive because only a few samples were analyzed due to budget constraints. The goal was just to find presence of isotope in non-labeled seedlings to demonstrate the existence of a fungal network connecting both seedlings at the start of the experiment. A synthesized version of this figure is provided in Figure 3 of the main manuscript.

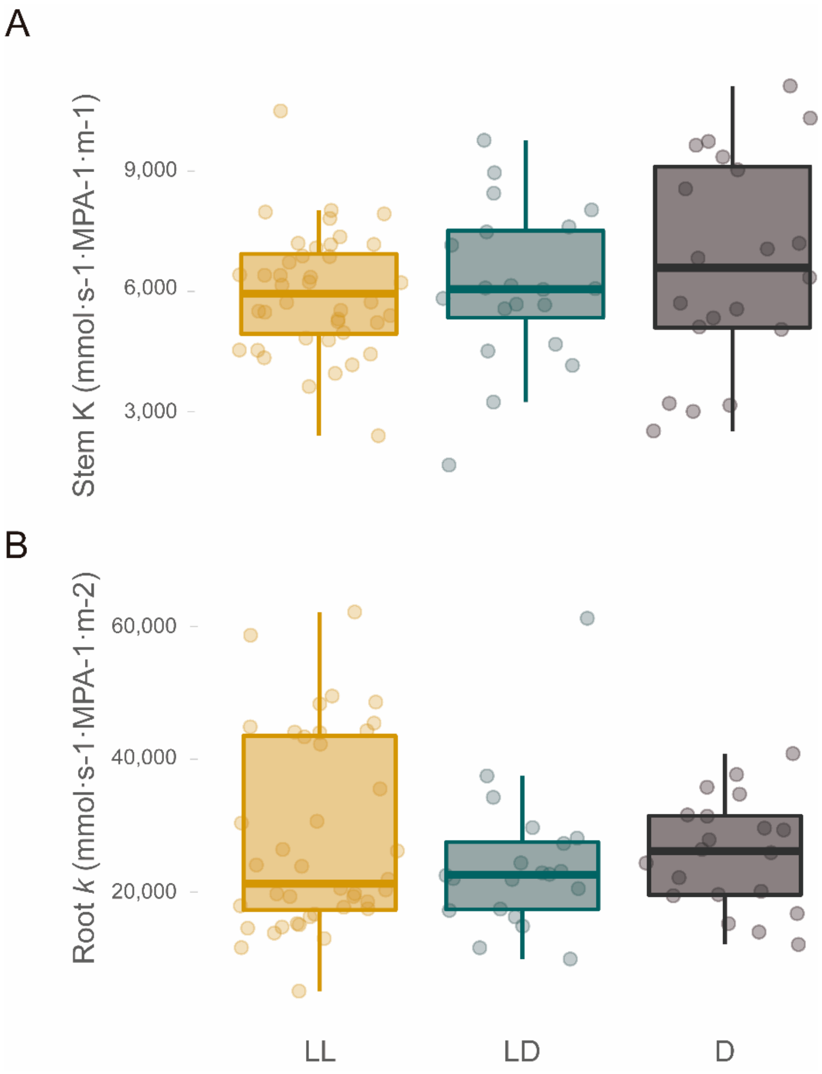


**Fig. S9.** Stems and roots of NSC-depleted plants did not lose water transport capacity. Panel A corresponds to hydraulic conductivity in stems. Panel B corresponds to hydraulic conductance in roots. Colors represent LL seedlings exposed to natural light (golden), D seedlings with light-blocking covers (black), and LD seedlings exposed to natural light paired with plants with covers (teal). Lines within boxes represent the median and top and bottom hinges represent 25th and 75th percentiles. Whiskers indicate highest and lowest value no further than 1.5 times the inter-quartile range represented by the hinges. Dots represent the distribution of the data. Asterisks indicate the degree of significance between groups (* = 0.05, ** = 0.01, *** = < 0.001).

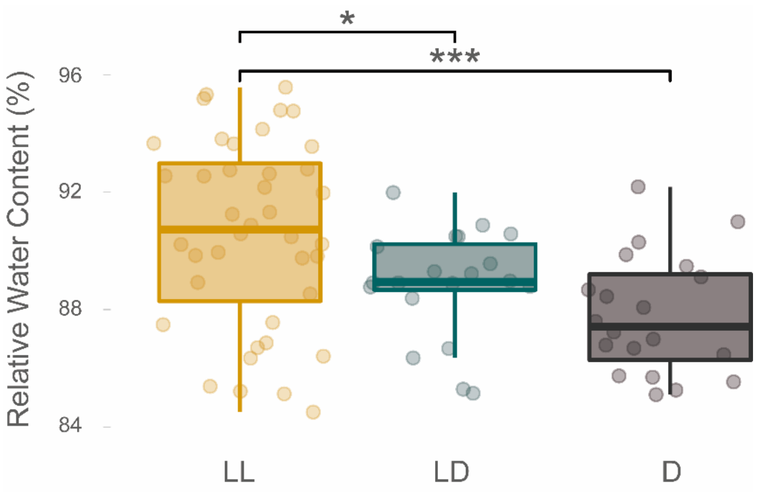


**Fig. S10.** NSC-depleted plants lose stem water content under well-watered conditions. Colors represent LL seedlings exposed to natural light (golden), D seedlings with light-blocking covers (black), and LD seedlings exposed to natural light paired with plants with covers (teal). Lines within boxes represent the median and top and bottom hinges represent 25th and 75th percentiles. Whiskers indicate highest and lowest value no further than 1.5 times the inter-quartile range represented by the hinges. Dots represent the distribution of the data. Asterisks indicate the degree of significance between groups (* = 0.05, ** = 0.01, *** = < 0.001).

**Table S2.** Linear models showing the influence of non-structural carbohydrate depletion on pressure and osmotic potentials. A constant was added to Leaf NSC deviation from control that would allow a log transformation necessary to meet model assumptions by shifting all values to a positive range.

| Model and Factors | Estimate | 95% C.I. Estimates | | *p-value* | d.f. (res.) | R | Adjusted R square |
|---|---|---|---|---|---|---|---|
| | | 2.5% | 97.5% | | | | |
| *Leaf Osmotic Potential = log(Leaf NSC Deviation from Control+100)* | | | | **<0.001** | 75 | -0.83 | 0.69 |
| Intercept | 1.93823 | 1.366727 | 2.5097296 | **<0.001** | - | - | - |
| *log(Leaf NSC Deviation from Control+100)* | -0.89378 | -1.030518 | -0.7570468 | **<0.001** | - | - | - |
| *Leaf Pressure Potential = log(Leaf NSC Deviation from Control +100)* | | | | **<0.001** | 75 | 0.64 | 0.40 |
| Intercept | -2.0313 | -2.8663738 | -1.1961497 | **<0.001** | - | - | - |
| *log(Leaf NSC Deviation from Control+100)* | 0.7326 | 0.5328177 | 0.9324307 | **<0.001** | - | - | - |
| *Leaf Pressure Potential = Leaf Water Potential x Plant NSC Deviation from Control* | | | | **<0.001** | 71 | 0.78 | 0.60 |
| Intercept | 2.444733 | 2.038011306 | 2.85145512 | **<0.001** | - | - | - |
| *Leaf Water Potential* | 1.426828 | 0.936520416 | 1.91713622 | **<0.001** | - | - | - |
| *Plant NSC Deviation from Control* | 0.022895 | 0.014616741 | 0.03117281 | **<0.001** | - | - | - |
| *Leaf Water Potential x Plant NSC Deviation from Control* | 0.014489 | 0.004241656 | 0.02473690 | **0.006** | - | - | - |
